# Integrating Mendelian randomization and multiple-trait colocalization to uncover cell-specific inflammatory drivers of autoimmune and atopic disease

**DOI:** 10.1101/394296

**Authors:** Lucy M. McGowan, George Davey Smith, Tom R. Gaunt, Tom G. Richardson

## Abstract

Immune mediated diseases (IMDs) arise from a lack of immune tolerance, causing chronic inflammation. Despite their growing prevalence, targeted therapies to treat IMDs are lacking. Cytokines and their receptors, which mediate inflammation, have been associated with IMD susceptibility. However, the complex signalling networks and multiple cell-types required to orchestrate inflammatory responses have made it difficult to pinpoint specific cytokines and immune cell-types which drive IMDs.

In this study, we developed an analytical framework which integrates Mendelian randomisation (MR) and multiple-trait colocalization (moloc) analyses to determine putative cell-specific drivers of IMDs. We used MR to determine the likelihood of causal associations between the levels of 10 circulating cytokines/cytokine receptors and 9 IMDs within human populations of European descent. Conservative (single SNP) and liberal (multiple SNP) MR analysis supported a causal role for IL-18 in inflammatory bowel disease (P = 1.17 × 10^−4^) and eczema/dermatitis (P = 2.81 × 10^−3^), as well as roles for IL-2rα and IL-6R in several IMDs.

Where associations between cytokines/cytokine receptors and IMDs were discovered using MR, we undertook moloc analyses. This was to assess the likelihood that cytokine/cytokine receptor protein quantitative trait loci (pQTL) and IMD-associated loci share a causal genetic variant along with expression QTL (eQTL) using data from 3 immune cell-types: monocytes, neutrophils and T cells. We found a monocyte and neutrophil-driven role for IL-18 in IBD pathogenesis, amongst evidence supporting several other cell-specific inflammatory drivers of IMDs. Our study helps to elucidate causal pathways for the pathogeneses of IMDs which, together with other studies, highlights possible therapeutic targets for their treatment.

## Introduction

Autoimmune and atopic diseases may arise due to a lack of immune tolerance towards self-antigen or harmless allergens respectively^1^. Loss of immune tolerance results in recurrent or chronic inflammation, causing damage to healthy tissues and extensive morbidity. The incidence of immune-mediated diseases (IMDs) has drastically increased in recent decades, highlighting the need for a clearer understanding of their pathogeneses and effective drug discovery^2^. Cytokines and growth factors (herein referred to as cytokines) are signalling factors which orchestrate the balance between immune homeostasis and inflammation^3^. However, traditional observational epidemiological studies are prone to confounding and reverse causation, making it challenging to disentangle causal effects of individual cytokines on IMDs^4^.

Genome-wide association studies (GWAS) have been instrumental in identifying large numbers of genetic loci which influence disease risk. This includes associations between loci which harbour genes responsible for the synthesis of cytokines and their receptors with autoimmune diseases such as inflammatory bowel disease (IBD)^5-8^, multiple sclerosis (MS)^9-11^, rheumatoid arthritis (RA)^12^ and systemic lupus erythematosus (SLE)^13^, as well as atopic diseases such as eczema^14^ and asthma^15^. This suggests that particular inflammatory cytokines may have a causal effect on the development of these diseases^4^. Previous studies have not yet integrated genome-wide association and cytokine loci data with cell or tissue-specific gene expression loci data to characterise the molecular basis of IMD pathogenesis. Identifying immune-cell specific disease-drivers, as well as putative causal relationships between cytokines and IMDs, will help to elucidate complex IMD pathways and identify drug target candidates for therapeutic intervention. Furthermore, targets supported by evidence from statistical analyses of human genetic data are thought to have double the success rate in clinical development^16^. Mendelian randomization (MR) is an increasingly popular statistical method used to strengthen causal inference with respect to exposure-disease associations within human populations, in the absence of confounding variables. MR uses single nucleotide polymorphisms (SNPs), identified through GWAS, as unconfounded proxies for an exposure of interest, analogous to a randomized controlled trial^17^. In this study, we used a conservative (single SNP) and liberal (multiple SNPs) two-sample MR analysis to investigate associations between 10 circulating inflammatory biomarkers (cytokines or cytokine receptors^18-23^) (Table S1) and 9 IMDs (Table S2). We subsequently used the recently-developed multiple-trait colocalization (moloc) method^24^, integrating either immune cell or tissue expression quantitative expression loci (eQTL), cytokine protein QTL (pQTL) and IMD-associated loci data, to identify immune cell and tissue-specific drivers of inflammatory disease.

## Methods and Materials

### Data Sources

For our two-sample MR analysis, we harnessed genetic instrument data for 10 circulating inflammatory cytokines or soluble cytokine receptors (Table 1) from summary statistics of previously published GWAS^18-20; 25^. The SNP used as an instrument for *IL-6R* affected levels of soluble IL-6R. The SNPs chosen in this analysis have been shown to be robustly associated with a change in circulating levels of a cytokine (P < 5 × 10^−8^) and are in *cis* with the gene of interest (i.e. the SNP was located within a 1MB distance of the gene which encoded the cytokine or cytokine receptor of interest). Data concerning IMD outcomes (Table 2) were derived from large-scale GWAS using the MR-Base platform^26^. For the moloc analysis, we harnessed human monocyte (CD14^+^ CD16^−^), neutrophil (CD16^+^ CD66b^+^) and T cell (CD4^+^ CD45RA^+^) eQTL data from the BLUEPRINT epigenome project^27^. All data used were derived from populations of European descent.

### Mendelian Randomization

Mendelian randomization follows 3 assumptions; that the selected instruments used are robustly associated with the exposure (1), that the selected instruments are unconfounded (2), and that the selected instruments can only influence the outcome via the exposure (3). Using randomly-inherited unmodifiable SNPs associated with circulating inflammatory cytokine levels through GWAS as genetic instruments for MR satisfies assumptions 1 and 2. We performed two-sample MR with the MR-Base platform^26^, using two different analysis methods (Figure S1A) as described below, depending upon available data:

Conservative two-sample MR was used to analyse the causal effect of cytokines using SNPs at target genes encoding the inflammatory biomarkers of interest. As such, for this analysis we used a single SNP acting in *cis* as a genetic instrument (i.e. located within a 1MB distance of the target gene encoding the cytokine with P < 5 × 10^−08^). As only one genetic instrument was used in this analysis, effect estimates were based on the Wald ratio test:

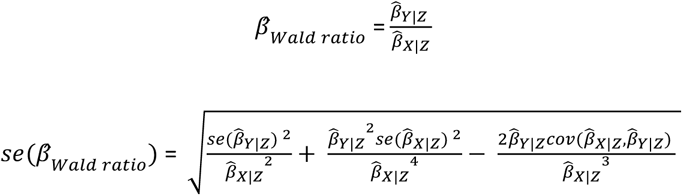

where 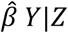 is the coefficient of the genetic variant in the regression of the exposure (e.g. circulating cytokine level) and 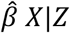 is the coefficient of the genetic variant in the regression of the outcome.

Where possible, liberal MR was also used to analyse the causal effect of circulating inflammatory biomarkers on immune-mediated diseases. In contrast to the conservative MR analysis, liberal MR used multiple SNPs as genetic instruments which were either acting in *cis* or *trans* (i.e. over 1MB distance from the target gene encoding the cytokine with P < 5 × 10^−08^). To identify instruments for the liberal MR we undertook genome-wide LD clumping based on P < 5 × 10^−08^ and r^2^ < 0.001. A leave-one-out MR analysis was performed in parallel with liberal MR to ensure that causal effects were not observed due to the influence of a single SNP. As two or more genetic instruments were available for liberal MR, we used the inverse variance weighted (IVW) method to obtain MR effect estimates:

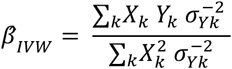

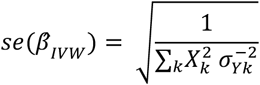

Where X is the exposure, Y is the outcome and the genetic variants are k (where k = 1,…,n).

For both conservative and liberal MR analyses, we used the Bonferroni correction to calculate an adjusted P value threshold (conservative: P ≤ 6.17 × 10^−4^, liberal: P ≤ 6.94 × 10^−4^) to account for multiple testing, where:

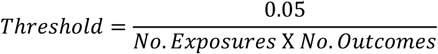

We subsequently undertook reverse MR analyses for all associations which survived multiple testing to investigate reverse causation (i.e. the likelihood that genetic predisposition to disease has an influence on inflammatory cytokine levels). As an additional sensitivity analysis, we used the MR Steiger directionality test to assess the directionality of associations between inflammatory cytokines and complex traits^28^.

### Multiple Trait Colocalization Analysis

We applied the moloc approach^24^ to immune cell-specific expression quantitative trait loci (eQTL) data, inflammatory biomarker protein QTL (pQTL) data, and IMD-associated loci data, to identify putative immune cell-specific drivers of IMDs (Figure S1B). The moloc method uses a Bayesian statistical framework to calculate PPA (posterior probability of association) scores to measure the degree of colocalization between gene loci using 3 or more datasets. PPA scores of ≥ 80% are likely to share a common genetic causal variant based on evaluations undertaken by the authors^24^. We obtained eQTL data derived from concerning neutrophils (CD16^+^ CD66b^+^), monocytes (CD14^+^ CD16^−^), and T cells (CD4^+^ CD45RA^+^)^27^. We ran independent analyses for each of the 3 immune cell-types, testing the degree of colocalization between immune cell eQTL, with inflammatory biomarker pQTL and IMD-associated loci.

We chose to perform moloc for inflammatory biomarkers and IMDs in this instance based upon a P-value threshold < 0.05 in either the conservative or liberal MR analysis. Along with investigating cell-type specificity for identified associations, this analysis was used to detect evidence of a coordinated system which is consistent with causality (i.e. gene expression and respective protein products colocalize with the associated complex traits). As such these findings can complement evidence from MR to detect putative causal effects between biomarkers and disease.

Finally, we undertook moloc analyses to investigate gene expression within different tissue types using data from the GTEx consortium v6p^29^. Our 3 traits in each analysis were the circulating inflammatory cytokine using pQTL data, the complex trait with strongest evidence of association with the cytokine in either the liberal or conservative MR analyses, and tissue-specific gene expression for the cytokine using GTEx eQTL data. We only investigated tissue-types with at least one eQTL (P < 1.0 × 10^−04^) for the target cytokine gene. As in the immune-cell eQTL moloc analysis, evidence of colocalization was based on a PPA score of ≥ 80%.

All statistical and bioinformatics analyses were undertaken using R statistical software version 3.31 ^30^. Plots illustrating multiple trait colocalization were generated using base R graphics, whereas our volcano plot was generated using ggplot^31^.

## Results

### Mendelian Randomization Analyses Identify Putative Causal Relationships Between Circulating Cytokine/Cytokine Receptor Levels and IMDs

We first used a conservative MR approach to detect associations between inflammatory biomarkers and IMDs, using a single cis-acting SNP instrument for the inflammatory biomarker gene of interest. *IP-10* and *TRAIL* were removed from the analysis as the SNP identified as instruments^18^ had minor allele frequencies too rare (< 0.05) to undertake formal two-sample MR. Results from the conservative MR analysis are displayed in Figure 1 and Table S3. Based on a Bonferroni corrected P-value threshold of P ≤ 6.17 × 10^−4^, we identified associations between soluble IL-6R levels and eczema/dermatits (P = 1.35 × 10^−8^), rheumatoid arthritis (RA) (P = 5.67 × 10^−8^), Crohn’s disease (CD) (P = 2.81 × 10^−5^) and Asthma (P = 1.12 × 10^−4^), as well as an association between IL-2Rα levels and multiple sclerosis (MS) (P = 1.75 × 10^−5^).

**Figure 1.**
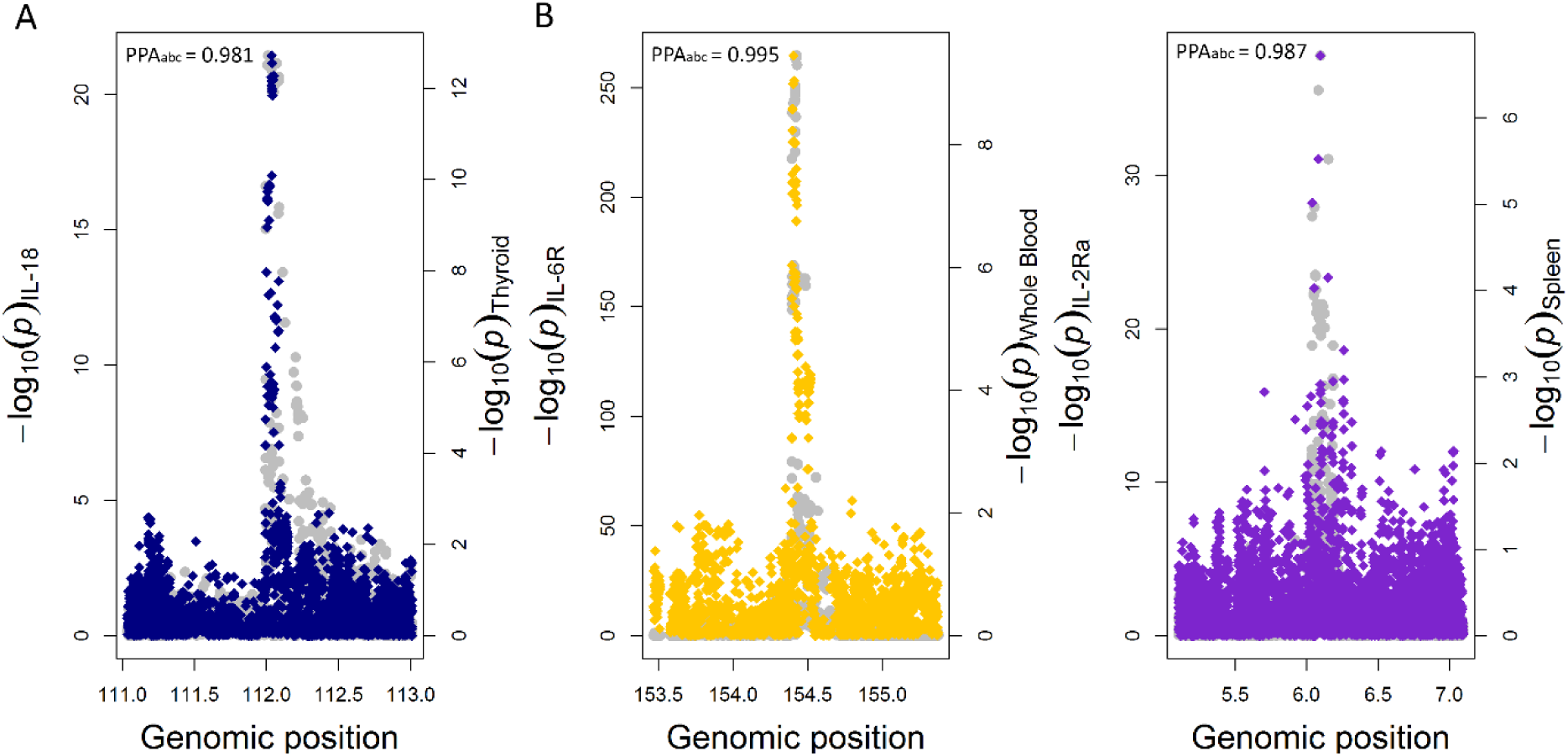
Conservative Mendelian randomization (MR) analysis detects associations between circulating inflammatory biomarkers and IMDs. Volcano plot of conservative MR analysis illustrating associations between inflammatory cytokines and complex traits. Red (upper) line represents Bonferroni corrected threshold (P ≤ 6.17 × 10^−4^) and black dotted line (bottom) represents unadjusted threshold (P ≤ 0.05).

We next performed a liberal MR analysis using all available instruments (i.e. acting in either *cis* or *trans*), associated with our inflammatory biomarkers (Exposure, Table S1) as instruments. *IL-6R* (SNPs affecting levels of soluble IL-6R only), *MIF*, and *IL-2Rα* were excluded from the liberal analysis as there was only one remaining SNP available for MR after LD clumping to remove SNPs in linkage disequilibrium with one another. We adjusted our Bonferroni corrected P value threshold to account for the change in the number of tests and applied the new threshold of P ≤ 6.14 × 10^−4^ to the liberal MR results. From this analysis we found evidence to suggest there may be causal relationships between circulating levels of IL-18 and inflammatory bowel disease (IBD) (P = 1.17 × 10^−4^) and eczema/dermatitis (P = 2.81 × 10^−3^). These associations were not detected in the conservative analysis using the *IL-18* SNP (rs71478720) alone (P = 1.06 × 10^−2^, E/D: P = 2.81 × 10^−3^), although using multiple instruments in the liberal MR provided much stronger evidence of association.

One of the advantages of the liberal MR analysis is that sensitivity analyses can be performed to test the robustness and the direction of putative inferred causal relationships. Thus, we next conducted a leave one out analysis to ensure that no single SNP from the instruments was responsible for the observed effect. Both the analyses of *IL-18* on IBD (Figure 2) and on eczema/dermatitis (Figure S2) survived leave-one-out analyses (Table S5), as the removal of any individual SNP from the analysis had little effect on observed effect estimates. These results provide evidence to support a causal role for circulating IL-18 levels in the pathogenesis of IBD and eczema/dermatitis. Moreover, reverse MR (Table S6) and the Steiger directionality test^28^ (Table S7) showed that reverse causation was unlikely for any of the associations identified in either the conservative or liberal MR analysis.

**Figure 2.**
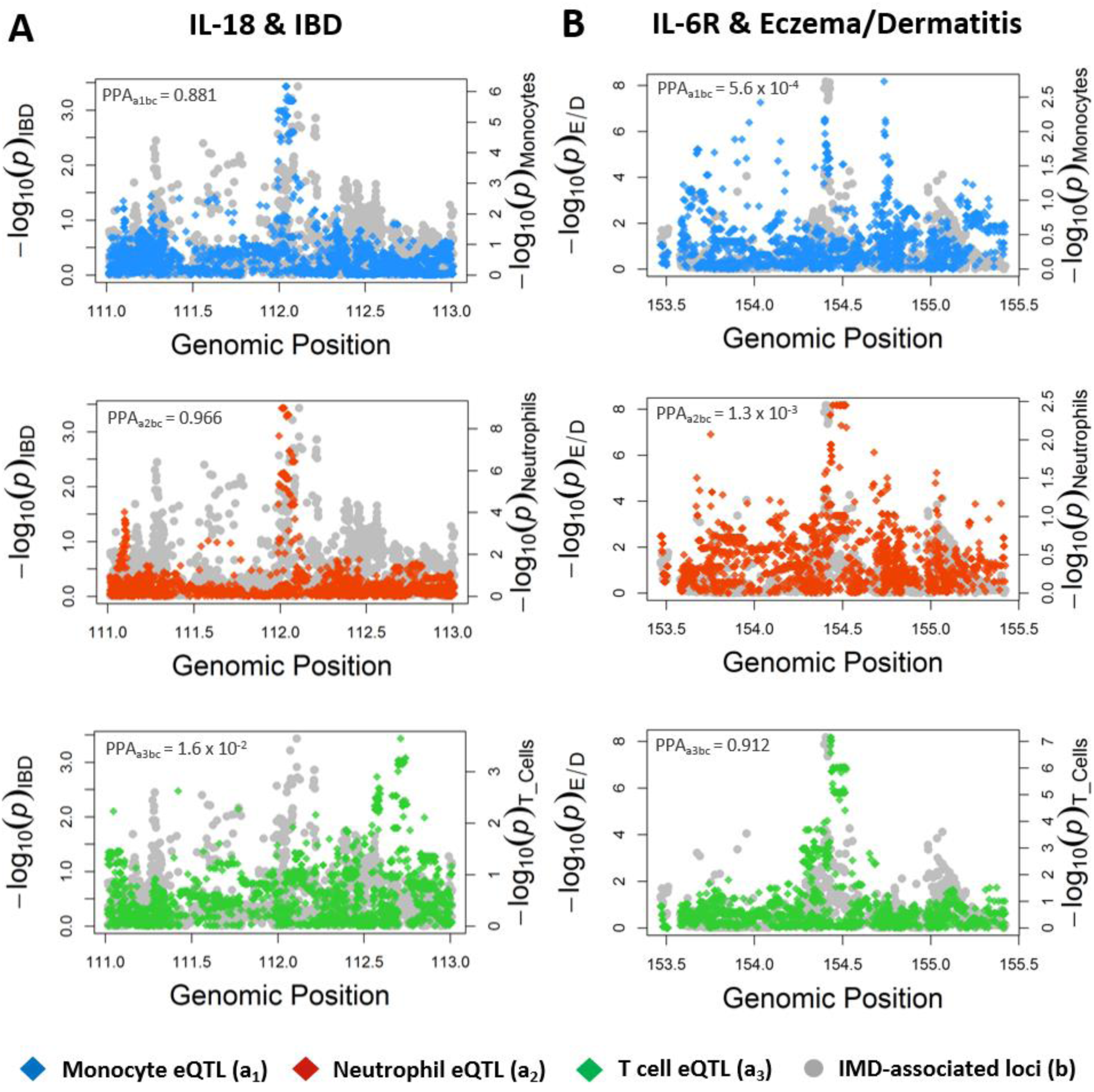
Liberal Mendelian randomization (MR) identifies a putative causal relationship between circulating levels of IL-18 and IBD which survives leave-one out sensitivity analysis. Leave-one out MR analysis for SNPs used as instruments for liberal MR analysis (black). Results show that this effect is not likely to be due to an individual SNP when compared to the observed effect of all SNPs (red).

### Multiple-trait Colocalization Uncovers Immune Cell-Specific Drivers of IMD

We next investigated the cell-type specificity for the detected putative drivers of IMD using eQTL data from the BLUEPRINT consortium^27^. For circulating inflammatory biomarkers and IMDs where evidence of a causal association was detected using MR, we applied the moloc method using eQTL data for the gene encoding the target inflammatory biomarker, pQTL data for the circulating inflammatory biomarker itself and GWAS summary statistics for the associated complex trait (Figure S1B). We tested whether immune-cell eQTL (_a_) colocalised with pQTL (_b_) and IMD-associated loci (_c_) The moloc analysis was performed 3 times for each association, using eQTL from either human CD14^+^ CD16^−^ monocytes (_a1_), CD16^+^ CD66b^+^ neutrophils (_a2_) or CD4^+^ CD45RA^+^ T cells (_a3_).

We found evidence of multiple-trait colocalization between 10 combinations of immune-cell eQTL, IMD-associated loci, and inflammatory biomarker pQTL (Table S8). PPA scores measuring colocalization of genetic signals between all 3 traits (PPA_abc_) indicated that monocytes (PPA_a1bc_ = 0.8812) and neutrophils (PPA_a2bc_ = 0.9657), but not T cells (PPA_a3bc_ = 0.0159), may drive IBD via IL-18 (Figure 3A). Additionally, we found evidence of a T cell-specific role in the disease pathways of eczema/dermatitis (PPA_a3bc_ = 0.9115) driven by soluble IL-6R (Figure 3B). IL-2Rα pQTL, MS-associated loci and T cell eQTL showed evidence of colocalization (PPA_a3bc_ = 0.8826), but there were no *IL-2Rα* eQTL data within monocytes or neutrophils in the BLUEPRINT study.

**Figure 3.**
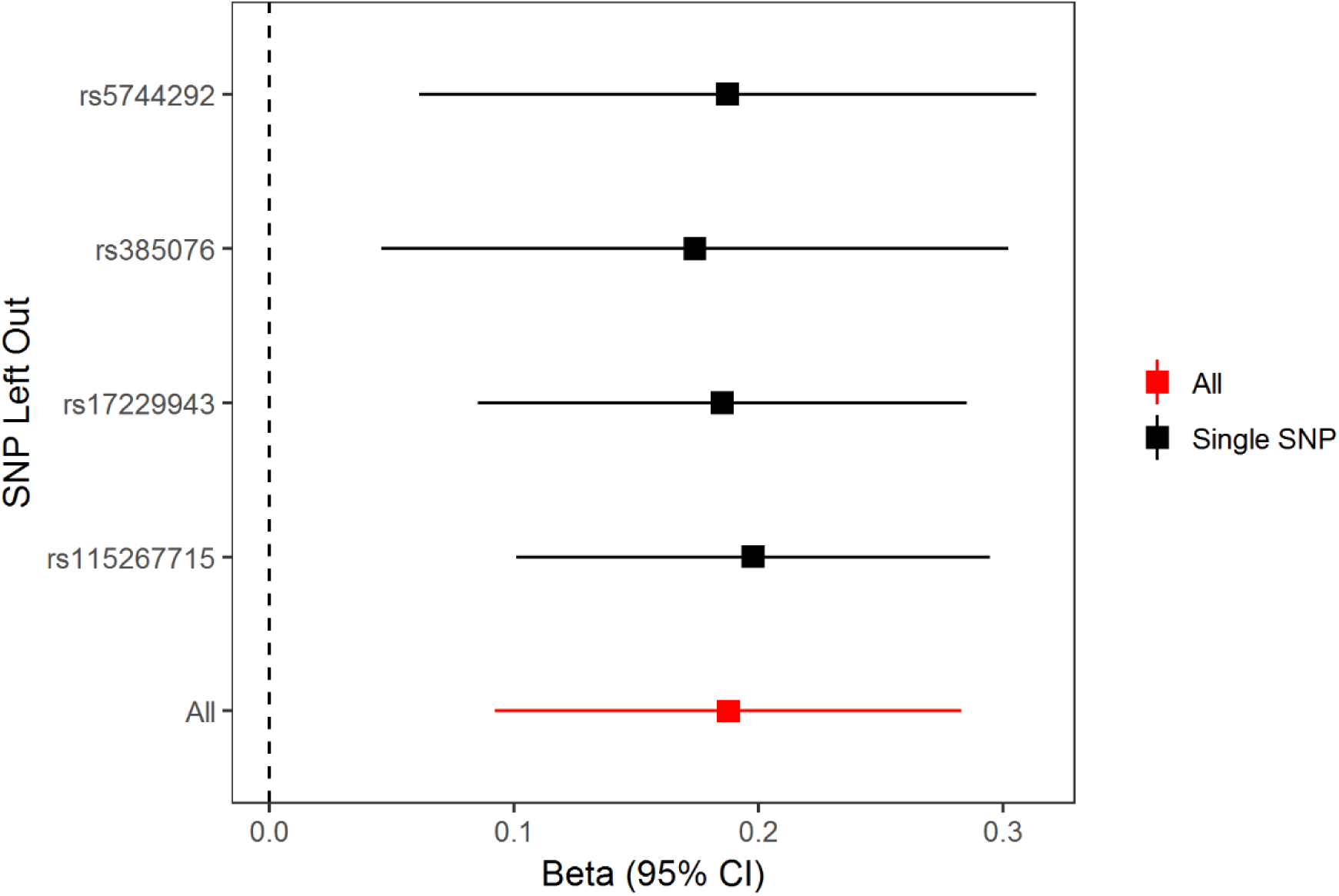
Multiple-trait colocalization reveals immune cell-specific divers of IMDs using immune cell expression quantitative trait loci (eQTL) data, inflammatory cytokine or cytokine receptor protein QTL (pQTL) data and immune-mediated disease GWAS data. These plots illustrate observed effects of genetic variants at the *IL-18* (A) and *IL-6R* (B) loci on inflammatory bowel disease (IBD) and eczema/dermatitis (E/D) respectively. Effect estimates on the expression of IL-18 and IL6R are overlaid in each plot using eQTL data derived from monocytes (top, _a1_), neutrophils (middle, _a2_) and T cells (bottom, _a3_). For simplicity, circulating cytokine effects are not displayed within the plots but were used to calculate PPA_abx_ scores. PPA_abc_ values reflect the likelihood that a causal variant influences the target cytokine (_b_), associated complex trait (_c_) and the expression of the corresponding gene (_a_). PPA_abc_≥ 0.8 indicates evidence of colocalization (i.e. a shared genetic variant between all 3 signals) and suggests that the cytokine (or its receptor) is a putative driver of the IMD when it is expressed within the cell-type of interest.

In the tissue-specific analysis, the association between IL18 and IBD was relatively ubiquitous, as evidence of colocalization was observed within 7 diverse tissue types (Table S9), including thyroid tissue (Figure 4a). Evidence of colocalization for the association between soluble IL6R and eczema/dermatitis was observed in 3 tissue types, most strongly in whole blood (PPA = 99.5%), which may help shed light on the pleiotropic effects observed at this locus (Figure 4b). Lastly, the association between IL2ra and multiple sclerosis colocalized in only two tissues; subcutaneous adipose and spleen (Figure 4c).

**Figure 4.**
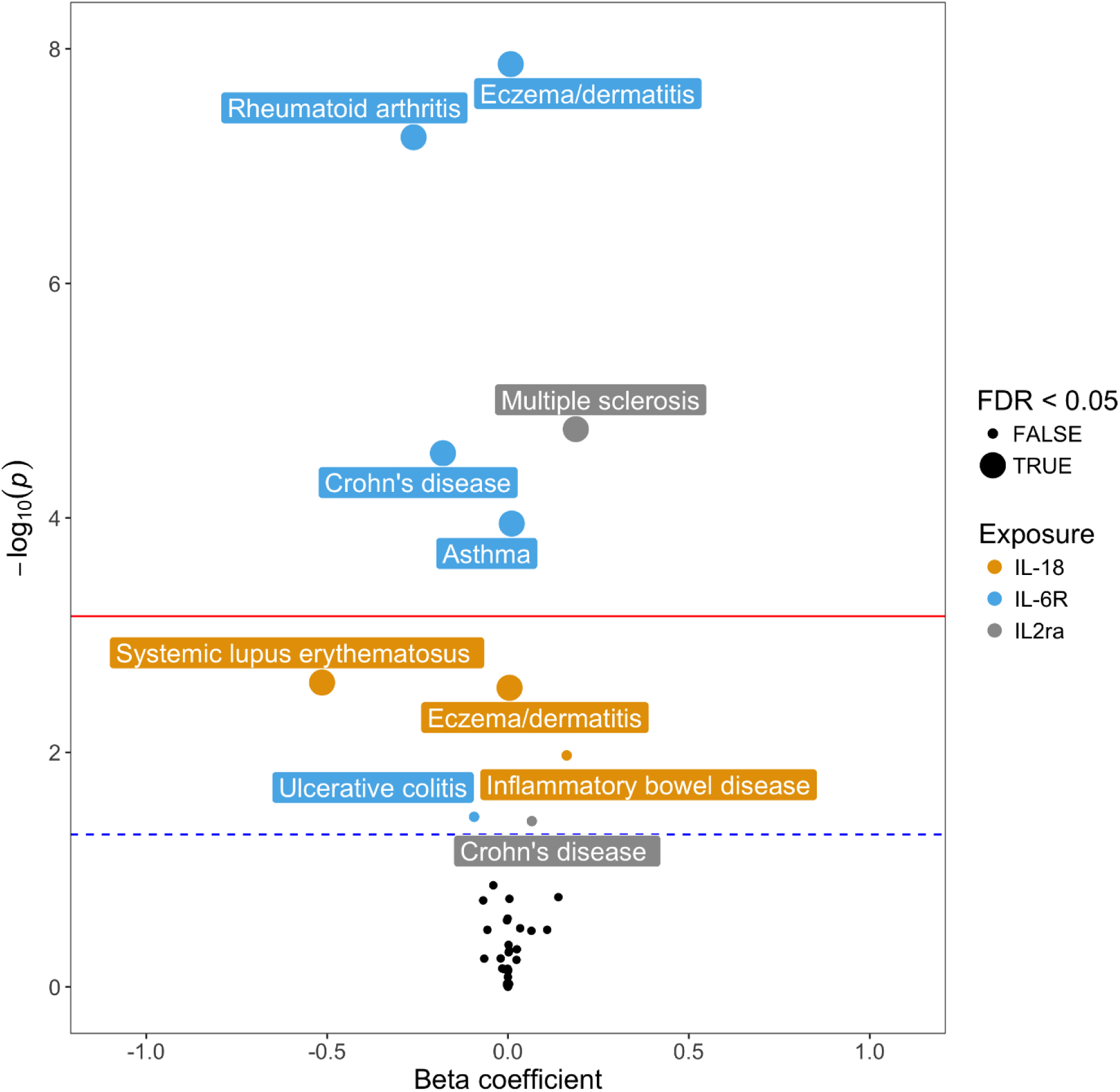
Multiple-trait colocalization indicates tissue specific expression of genes which are associated with circulating cytokine or cytokine receptor levels and immune-mediated disease (IMD) susceptibility. These plots illustrate observed effects of genetic variants at the *IL-18* (A), and *IL-6R* (B) and *IL-2Rα* (C) loci with their corresponding protein product. Effect estimates on the expression of IL-18, IL-6R, IL-2Rα are overlaid in each plot using expression quantitative trait loci data derived from thyroid (A), whole blood (B) and spleen (C). For simplicity, effects on complex traits are not displayed within the plots but were used to calculate PPA_abx_ scores. PPA_abc_ values reflect the likelihood that a causal variant influences the target cytokine (_b_), associated complex trait (_c_) and the expression of the corresponding gene (_a_). PPA_abc_ ≥ 0.8 indicates evidence of colocalization (i.e. a shared genetic variant between all 3 signals) and suggests that the cytokine (or its receptor) is a putative driver of the IMD when it is expressed within the tissue-type of interest.

## Discussion

The prevalence of autoimmune and atopic diseases has increased drastically over recent decades, particularly within European populations^2; 32^. Despite the rising demand, a poor understanding of the molecular pathways driving IMDs means that drugs for their treatment are lacking^33; 34^. Cytokines and their receptors mediate the balance between tolerance and inflammation, by signalling between immune cells, making them attractive therapeutic targets^35^. However, the ways in which the hundreds of functionally diverse cytokines interact and orchestrate inflammatory responses in the body are unclear^36^. Consequently, identifying individual cytokines for drug-targeting which drive IMDs has been difficult. The use of two-sample MR has aided the elucidation of many disease pathways, by measuring the likelihood of causal relationships between exposures and traits in human populations^37^. In this study, we developed a framework that integrates MR with multiple-trait colocalization to gain insights into the molecular basis of IMD pathogenesis. Using this framework, we found evidence to support causal relationships between levels of: circulating IL-18 and IBD, as well as eczema/dermatitis, circulating soluble IL-6R and eczema/dermatitis, and circulating IL-2Rα and MS, amongst others (Tables S3-4, S6-8). Additionally, we provided evidence to suggest that these associations are likely to be driven in an immune-cell and tissue-specific manner. We believe our analysis framework could be applied by other studies with alternative hypotheses, as a way of disentangling complex biochemical cell signalling pathways and identifying molecules and cell-types which are likely to drive disease.

A T cell-mediated role for IL-2/IL-2Ra in MS has already been well established through epidemiological and lab-based studies^10; 38^. The IL-2R-targetting drug daclizumab was given FDA approval for the treatment of MS, but was recently withdrawn due to serious side-effects^39-41^. Additionally, a causal role for IL-18R in atopic dermatitis has recently been described, by integrating MR and pQTL-data^42^. Our results provide additional evidence to support these existing findings, as well as identifying monocytes and neutrophils as potential drivers of the relationship between IL-18 and eczema/dermatitis (Table S8). However, the associations between soluble IL-6R variation and eczema/dermatitis, or IL-18 and IBD within human populations are less well understood. Our analyses not only help establish evidence for causal relationships between these inflammatory biomarkers and IMDs, but also help characterize their cell-type specific nature.

IL-18, a member of the IL-1 superfamily of cytokines, is a potent inducer of Th1 mediated inflammation and INF-γ production^43^. This cytokine was first linked to IBD nearly 20 years ago, where it was shown to be highly expressed in intestinal tissues derived from IBD patients, compared to healthy control patients^44^. Using murine IBD models, deletion of *il-18* or its receptor *il-18r1* has been shown to be protective against inducible-colitis, by controlling goblet cell function and maintaining intestinal barrier homeostasis^45^. Despite strong evidence to suggest a role for IL-18 in IBD in mice, whether there was a causative role within human populations remained unclear. Our study provides evidence that *IL-18* is likely the causal gene responsible for the association with IBD at this locus, as well as out MR analysis supporting a causal role for IL-18 in the disease pathogenesis of IBD within human populations. Furthermore, the moloc analyses suggested that innate immune cells such as monocytes and neutrophils are likely to drive this association, supporting the current dogma that innate production of IL-18 stimulates Th1/Th17-mediated autoimmunity in IBD^47^. The T cell-specific eQTL data currently available for moloc analyses concerned naïve CD4^+^ T cells^27^. More eQTL data concerning activated and differentiated immune cell subsets, such as macrophages, dendritic cells and T helper cell subsets (i.e. Th1, Th2, Th17, Treg) are required for additional immune cell subset-specific moloc analysis, to further elucidate the molecular pathways which drive IMDs. However, this data may be difficult to acquire on a large-scale. Although eQTL data for many tissue types are now readily available^29^, moloc analysis using tissue-specific eQTL data concerning tissues which are known to become inflamed during IMDs would help to indicate whether cytokine or cytokine receptor expression in certain tissue types drives IMDs. For example, eQTL data from the different layers within the intestine would help to further unravel which roles inflammatory biomarkers are likely play in driving inflammation during IBD, in a tissue-context-dependent manner.

IL-6R is the receptor of the pro-inflammatory cytokine IL-6, which can exist in a membrane-bound state (classical) to the surface of leukocytes and hepatocytes, or in soluble form (trans)^48^. Both, classical and trans IL-6R signalling culminates in the expression of signal transducer and activator of transcription (STAT)3, which promotes inflammation via the expression of genes encoding antiapoptotic proteins and cytokines^49^. However, classical IL-6 signalling can act on few cell types compared to trans IL-6 signalling, which can act on any cell which has the cell-bound signal transducer, gp130^50^. One GWAS previously reported the an association between elevated soluble IL-6R levels resulting from a SNP in *IL-6R* and atopic dermatitis^51^. We provide evidence from the conservative MR and moloc analyses that supports a causal role between soluble IL-6R and eczema/dermatitis. Levels of membrane-bound IL-6R (leading to classical IL-6 signalling) and soluble IL-6R (trans IL-6 signalling) fluctuate as a result of receptor shedding by cells expressing membrane-bound IL-6R, as wells changes to the amount of receptor being synthesised by cells^50^. Moreover, the ratio of classical to trans IL-6 signalling also depends upon levels of IL-6 and cell-bound gp130. Therefore, analyses combining instruments which affect the classical or trans signalling pathways would provide a clearer insight into the role of IL-6 signalling in disease. Interestingly, activated CD4^+^ T cells have been shown increase levels of soluble IL-6R, by shedding their membrane-bound IL-6R; this mechanism is thought to have a role in the development of autoimmune diseases, which are often mediated by autoreactive T cells ^52; 53^. Through the moloc analysis, we showed that eczema/dermatitis was likely to be driven by IL-6R expression in T cells (Figure 3B, Table S8), a finding which is supported by evidence of increased IL-6R shedding leading to an increase in soluble IL-6R in people diagnosed with atopic dermatitis compared to healthy controls^51^.

Relatively few SNPs have been associated with changes in the circulating inflammatory biomarkers investigated in this study, as most of the GWAS summary statistics used to identify our instruments were derived from cohorts with fewer than 9000 people^18-20^; this is likely due to the high cost of quantification of circulating cytokines and cytokine receptors from blood. Our reverse MR analysis may therefore have been underpowered to evaluate evidence of reverse causation, although supplementing this analysis using the Steiger directionality test also suggested that this was unlikely for the associations we identified. As sample sizes for GWAS of circulating cytokines increase, the analysis pipeline illustrated by this study will have further power to detect novel relationships between markers of inflammation and complex disease.

In conclusion, in this study we have found strong evidence supporting new and known causal, immune-cell driven relationships between inflammatory biomarkers and IMDs. Triangulation of results from these analyses, with published results from experimental models of IMDs and GWAS, suggests that targeting IL-18 or its receptor, IL-18R, may be promising for the treatment of IBD. Indeed, small molecule inhibitors which target and repress IL-18-mediated signalling events are currently under development, although not for the treatment of IBD^54^. Moreover, a recently-published phase-II trial of an IL-18 binding protein (IL-18bp) drug to treat adult-onset Still’s disease demonstrated a favourable efficacy safety profile. If further trails are deemed successful, IL-18bp drugs may also be used to target other IMDs such as IBD.

## Appendices

No appendices.

## Supplemental Data Declaration

### Document S1

Supplementary figures S1 & S2 (PDF).

### Data S1

Supplementary tables S1-S8 (.xls).

## Acknowledgements

This study and L.M.M were financially supported by a 4-year studentship fund from the Wellcome Trust Dynamic Molecular Cell Biology Ph.D. Programme at the University of Bristol: (108907/Z/15/Z). T.G.R is a UKRI Innovation Research Fellow (MR/S003886/1). The research was carried out in the UK Medical Research Council Integrative Epidemiology Unit (MC_UU_00011/4 and MC_UU_00011/1). This study makes use of data generated by the BLUEPRINT Consortium^27^ and data generated by the GTEx Consortium^29^.

## Web Resources

MR-Base Platform: http://www.mrbase.org/

Blueprint: http://dcc.blueprint-epigenome.eu/#/home

GTEx: https://www.gtexportal.org/home/

